# Chemoproteomics reveals the epoxidase enzyme for the biosynthesis of camptothecin precursor strictosamide epoxide

**DOI:** 10.1101/2023.07.26.550496

**Authors:** Tong Zhang, Yan Wang, Shiwen Wu, Ernuo Tian, Chengshuai Yang, Zhihua Zhou, Xing Yan, Pingping Wang

**Affiliations:** CAS-Key Laboratory of Synthetic Biology, CAS Center for Excellence in Molecular Plant Sciences, Chinese Academy of Sciences, Shanghai, 200032, China; University of Chinese Academy of Sciences, Beijing, 100049, China

## Abstract

Camptothecin and its derivatives are the third largest anticancer drugs in the world market, mainly used to treat malignant tumors such as lung, colon and cervical cancer. Camptothecin was firstly discovered in *Camptotheca acuminate* and extracted mainly from *C. acuminate* and *Nothapodytes nimmoniana* for medicine production (Sadre et al. 2016). However, the overharvesting of *C. acuminate* and *N. nimmoniana* has greatly reduced their populations in nature, which are currently listed as the second protected plants in China and India. It is estimated there would be 20 million new cancer cases in 2025 all over the world, meeting the growing demand for camptothecin and other anti-cancer drugs has become a daunting challenge (Seca et al. 2018). In this study we tried to elucidate the unknown biosynthetic pathway from strictosamide **1** to strictosamide epoxide **2** by unearthing the candidate enzymes from the proteome of plant *Ophiorrhiza pumila* using the chemoproteomic strategy.

## Introduction

Due to the limited natural plant resources, high cost of chemical synthesis, producing camptothecin using synthetic biology method based on elucidation of synthetic pathway becomes a promising choice towards achieving this goal. However, the biosynthetic pathway of camptothecin has not yet been completely elucidated. In the camptothecin producing plant *Ophiorrhiza pumila*, the indole synthetic pathway providing the precusor tryptamine and the iridoid pathway providing the precusor secologanin have been elucidated. Product tryptamine and secologanin are catalyzed by strictosidine synthetase to form strictosidine, yet the downstream synthetic route from strictosidine to camptothecin remained unclear for more than a half century (Rai et al. 2021). It is speculated that strictosidine is firstly converted to strictosamide by reduction reaction, and then strictosamide was oxidized to strictosamide epoxide (Kang et al. 2021). After a series of oxidation, reduction and other reactions, strictosamide epoxide is converted to camptothecin (Fig S1).

The combinations of genomic, transcriptomic and proteomic analysis have usually applied to screen candidate genes and elucidate the unknown biosynthetic pathway of natural plant products (Wang et al. 2021). These omics data-based strategies have worked well in many cases, however, it often generate too many candidates, which may bring high labor-cost to screen target genes. Recently, chemoproteomic profiling based on the investigation of protein-small molecule interactions by combining functionalized chemical probes and proteomic analysis, has emerged as a promising tool to mine target enzymes in the biosynthetic pathways of natural products.

## Results and discussion

We tried to elucidate the unknown biosynthetic pathway from strictosamide **1** to strictosamide epoxide **2** by unearthing the candidate enzymes from the proteome of plant *Ophiorrhiza pumila* using the chemoproteomic strategy (Fig 1a). A bait for candidate protein fishing ST-probe **4**, with diazirine-alkyne tag, was designed and syntheiszed via 1,3-Dicyclohexylcarbodiimide promoting condensation of strictosamide **1** and 2-(3-(but-3-yn-1-yl)-3H-diazirin-3-yl) acetic acid (Dead-probe **3)** in anhydrous DMF. The structure of the synthetic probe was affirmed by HR-MS and 1D-NMR spectra.

**Figure 1.**
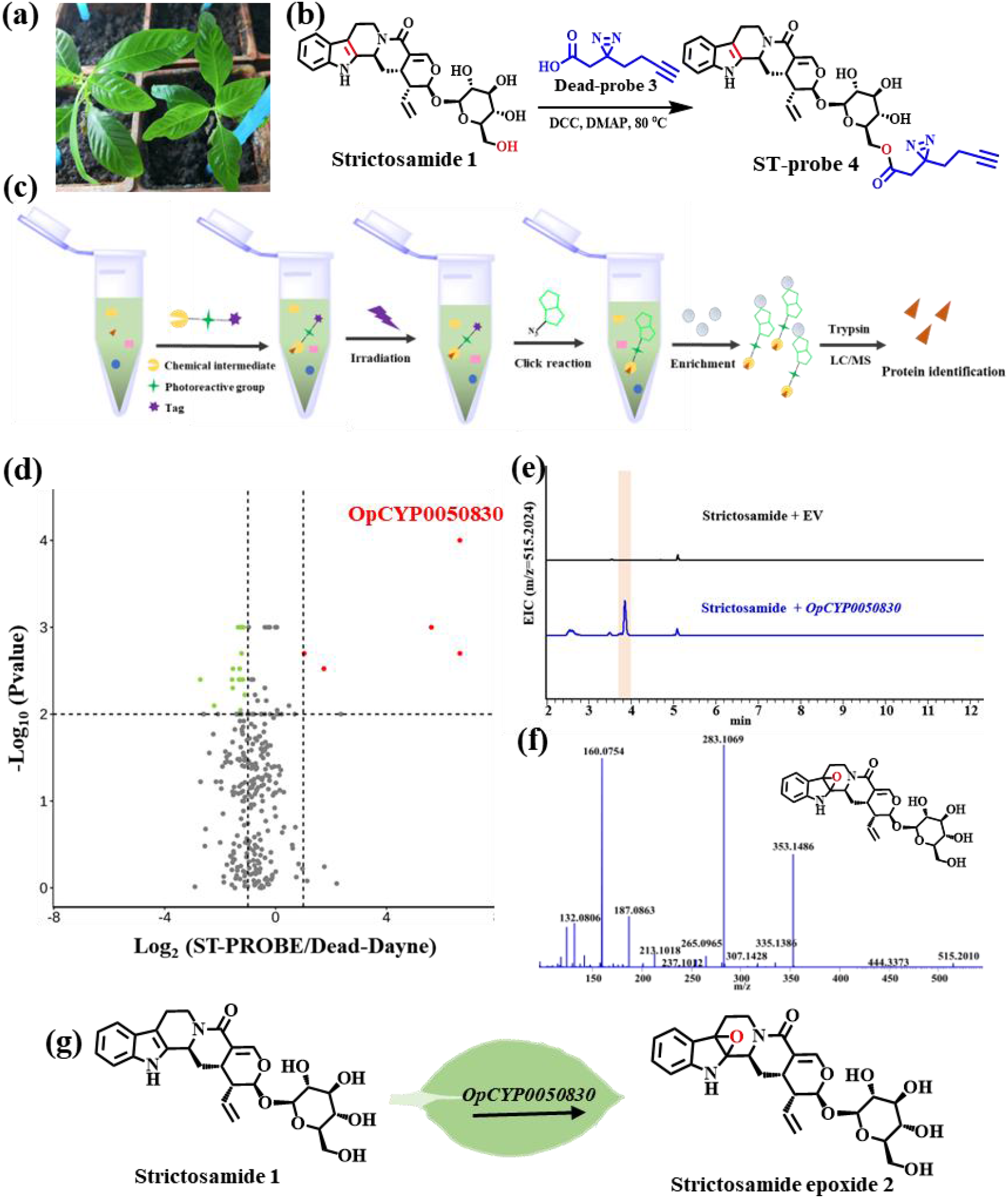
A CYP450 to catalyze strictosamide **1** forming strictosamide epoxide **2** has been identified by chemoproteomic analysis. (a) Image of a six-month-old *Ophiorrhiza pumila* plant. (b) Synthesis of probe ST-probe **4**. (c) Workflow of chemoproteomic profiling of *O. pumila* extract with ST-probe **4**. (d) Enrichment effect as shown by ST-probe **4**/Dead-probe **3**, blue point represents the increased proteins, green point represents the decreased proteins, grey point represents the proteins without change. (e) LC/MS analysis of the newly produced compound by *N. benthamiana* with transient expressed OpCYP0050830, extracted ion chromatograms for strictosamide epoxide **2** (m/z [M+H]^+^= 515.2024). (f) MS/MS spectra of strictosamide epoxide **2** produced in *N. benthamiana*. (g) The reaction catalyzed by OpCYP0050830.

To fish the corresponding protein which may catalyze strictosamide **1** to produce strictosamide epoxide **2**, the six-month-old *O. pumila* was harvested to extract the crude proteins (Fig 1b). The crude protein extract was then incubated with 100 μM ST-probe **4** and Dead-probe **3** (a negative control probe without the strictosamide motif) (Gao et al. 2020), respectively. The incubated proteins were then biotinylated, enriched and digested to be identified and relatively quantified by LC-MS/MS (Zhou et al. 2018), and 296 proteins were identified from the incubated *O. pumila* proteins with probe (Fig 1c). five proteins demonstrated more than 2 fold change (being considered as significant hits) in the ST-probe group in volcano plot (Table S1), of which a protein named as OpCYP0050830 was annotated as a CYP450(Fig 1d). The other four proteins were annotated as thaumatin-like protein, ATP synthase, superoxide dismutase and hypothetical protein, respectively, which are less likely to catalyze the epoxidation reactions. Therefore, OpCYP0050830 was further characterized as the candidate enzyme to produce strictosamide epoxide **2**.

To investigate the biochemical activity of OpCYP0050830, its corresponding gene OpCYP0050830 was cloned from the cDNA of an *O. pumila plant* and transformed into *Nicotiana benthamiana* for transient expression. After feeding strictosamide for 72 h, *N. benthamiana* leaves were extracted by methanol. A chromatographic peak with the same molecule weight of (M+H)^+^=515.2024 as the target compound strictosamide epoxide **2** could be detected in the *N. benthamiana* leave extract expressing OpCYP0050830, while not in leaves with transforming empty vectors (Fig 1e). The main ionic fragments of the newly produced compound, such as 515, 353, 335, 283, 265 and 237 were same with that of strictosamide epoxide **2** as previous reported (Sadre et al. 2016), which confirmed OpCYP0050830 could catalyze strictosamide **1** to produce strictosamide epoxide **2** (Fig 1f-g). Phylogenetic analysis revealed that OpCYP0050830 belong to the CYP716 family. The CYP716 family P450 PgCYP716S5 from *Platycodon grandiflorus* has previously been reported to catalyze the epoxidation of β-amyrin into 12,13α-epoxy β-amyrin (Miettinen et al. 2017).

There are estimated approximately 24, 000 genes by the annotation of *O. pumila* transcriptome, however, we could reduce the candidate number to 296 proteins based on chemoproteomics profiling analysis. Our results not only unveiled the long-time missing enzymes for the biosynthesis of strictosamide epoxide, but also providing an example showcase chemoproteomics as an efficiently tool for discovering unknown enzymes involved in natural products biosynthesis.

## Author contributions

Pingping Wang, Xing Yan, Zhihua Zhou and Tong Zhang designed the research. Tong Zhang performed the experiments, finished the sketch, analyzed the data and wrote the manuscript under the guidance of Pingping Wang and Zhihua Zhou. All authors contributed a lot to this work and approved the final version of the manuscript.

## Acknowledgements

This work was financially supported by the National Key Research and Development Program of China (Grant No. 2018YFA0900700), the National Natural Science Foundation of China (Nos. 31901021; 31921006), and the Strategic Biological Resources Service Network Plan of the Chinese Academy of Sciences (Grant No. KFJ-BRP-009). We thank Xiaoyan Xu and Shanshan Wang in the Core Facility Centre of CEMPS for help in analyzing chemical structures and protein sequences.

## Supporting information

**Fig S1** Supplementary Fifure.

**Table S1** Supplementary Table

