## Supplemental Figure 1 for "Chemoproteomics reveals the epoxidase enzyme for the biosynthesis of camptothecin precursor strictosamide epoxide"

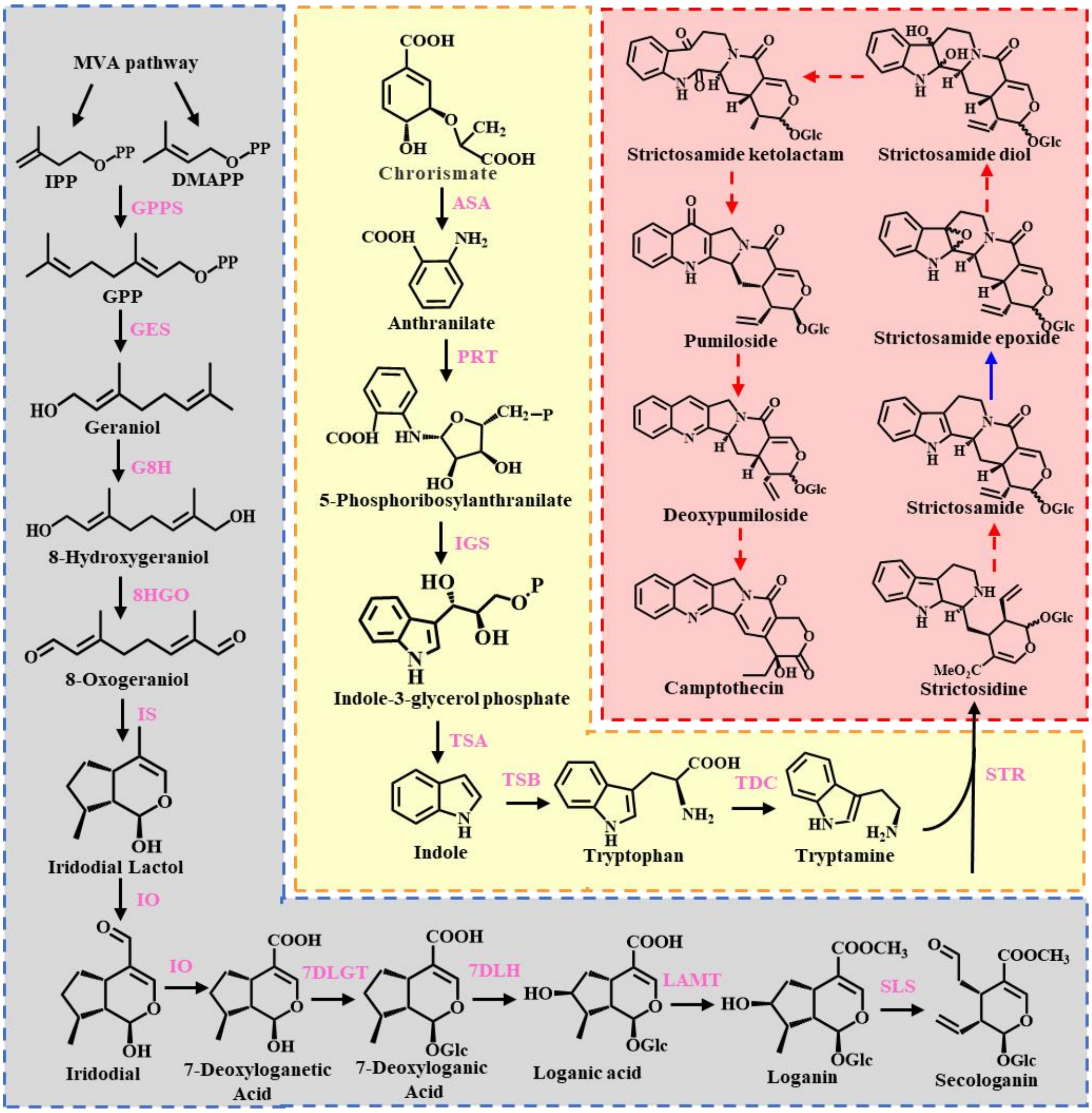


**Figure S1** The proposed biosynthetic pathway of camptothecin in *Ophiorrhiza pumila.* Black solid arrow represents previous identified reaction, red dotted arrow indicates an unknown pathway and blue solid arrow indicates identified reaction in this study.
